# Effects of acute adolescent stress on the acquisition and maintenance of intravenous oxycodone self-administration in male and female rats

**DOI:** 10.1101/2025.09.10.675431

**Authors:** Corinne A. Gallagher, Daniel J. Chandler, Daniel F. Manvich

**Affiliations:** Department of Neuroscience, Rowan-Virtua School of Osteopathic Medicine, 42 East Laurel Rd, Suite 2200, Stratford, NJ 08084; Program in Molecular Cell Biology and Neuroscience, Rowan-Virtua School of Translational Biomedical Engineering and Sciences, 42 East Laurel Rd, Suite 2200, Stratford, NJ 08084

**Keywords:** Prescription opioids, oxycodone, adolescent stress, self-administration

## Abstract

**Background:** The persistent threat of the opioid epidemic warrants investigation into risk factors that predispose individuals to opioid use disorder (OUD). Adolescent stress has been linked to enhanced risk for OUD in humans, however attempts to model this preclinically have yielded mixed results. Additionally, few studies have explored whether adolescent stress modulates the reinforcing effects of prescription opioids. Here we investigate the impact of acute adolescent stress on oxycodone self-administration in male and female rats.

**Methods:** Adolescent male and female rats underwent acute restraint stress during concurrent exposure to predator odor, or control handling. Approximately one week later, subjects were allowed to acquire IV oxycodone self-administration (0.03 mg/kg/inf) over 10 sessions (2 h/day) under a fixed-ratio 1 (FR1) schedule of reinforcement. Following three additional FR1 sessions and seven sessions under FR3, rats underwent two progressive-ratio tests (0.03 mg/kg/inf and 0.06 mg/kg/inf, respectively). Separate groups of adolescent rats underwent similar experimental manipulations but were trained on sucrose reinforcement.

**Results:** Adolescent stress did not affect the rate of acquisition of IV oxycodone self-administration. However, oxycodone self-administration escalated during post-acquisition FR1 sessions and remained elevated during FR3 sessions in stressed rats as compared to unstressed controls. Adolescent stress exposure did not affect responding during progressive-ratio tests, nor did it affect any measure of sucrose pellet reinforcement.

**Conclusions:** The present results are the first to demonstrate adolescent stress-induced enhancement of oxycodone reinforcement in rats and provide a preclinical model for investigating the neurobiological mechanisms by which adolescent stress increases vulnerability for prescription opioid misuse.

## 1. Introduction

The opioid epidemic is an increasing threat to public health, with opioid-involved overdose fatalities surpassing 81,000 in 2022 – a 64% increase from 2019 (National Institute on Drug Abuse, 2024). Initial opioid misuse often begins with prescription opioids (POs) prior to the development of opioid use disorder (OUD) (Dickson-Gomez et al., 2022; Jones, 2013; Lankenau et al., 2012; Veliz et al., 2022). PO misuse alone accounted for 88.2% of the 8.9 million self-reports of opioid misuse in people aged 12 or older in 2022 (Substance Abuse and Mental Health Services Administration, 2022). Moreover, an estimated 75% of young injection drug users report initiating opioid misuse with a PO (Lankenau et al., 2012). Among POs, oxycodone has been ranked by opioid-dependent individuals as the most desirable and “addictive”, and is the most commonly used PO by primary heroin users (Remillard et al., 2019; Rosenblum et al., 2007). Consequently, it is imperative to identify factors that may predispose individuals to oxycodone misuse in at-risk populations.

As a critical developmental period, the experience of stress or trauma during adolescence has been associated with increased risk for substance use disorders (SUD) (Basedow et al., 2020; Benjet et al., 2010; Dube et al., 2003; Scott et al., 2012; Skeer et al., 2009) as well as mood and/or anxiety disorders (e.g., major depressive disorder, bipolar disorder, generalized anxiety disorder, post-traumatic stress disorder) (Basedow et al., 2020; Benjet et al., 2010; Jaworska-Andryszewska and Rybakowski, 2019; McLaughlin et al., 2012; Scott et al., 2012) that in turn are linked with enhanced vulnerability for substance misuse and which often present comorbid with SUDs (Calarco and Lobo, 2021; Davis et al., 2017; Langdon et al., 2019; Patton et al., 2024; Preuss et al., 2021). In particular, OUD has been linked to a history of stress during adolescence (Austin and Shanahan, 2018; Conroy et al., 2009; Quinn et al., 2016; Swedo et al., 2020), with some estimates showing populations experiencing adolescent trauma have a 2.7-fold increase in opioid use (Heffernan et al., 2000). Despite these associations in clinical populations, preclinical studies investigating the effect of adolescent stress on opioid reinforcement in animal models are conflicting. For example, social isolation stress during adolescence increased self-administration of inhaled sufentanil (Weinhold et al., 1993) and IV remifentanil (Hofford et al., 2017) in male rats, and trended towards producing enhanced IV fentanyl self-administration in male and female rats (Bardo et al., 2023). However, other studies have reported no effect of early-life/adolescent social isolation stress on oral morphine consumption or IV remifentanil self-administration in adult rats (Marks-Kaufman and Lewis, 1984; Thorpe et al., 2020), while others have reported attenuated opioid-induced conditioned place preference following adolescent social isolation stress (Schenk et al., 1985; Wongwitdecha and Marsden, 1996). Notably, no studies to date have characterized whether oxycodone reinforcement specifically is affected by adolescent stress. Additionally, female subjects were not included in most prior studies, and all relied on social isolation as a stressor during adolescence.

The goal of this study was to examine whether acute adolescent stress impacts IV oxycodone self-administration in male and/or female rats. We employed a single exposure to predator odor during concurrent physical restraint as the stressor, based on previous evidence that presentation of this stressor during adolescence induces an anxiety-like behavioral and neurophysiological phenotype in male rats (Borodovitsyna et al., 2022, 2018). We first investigated whether exposure to this stressor during adolescence alters initial acquisition of oxycodone self-administration. We then assessed maintenance of oxycodone self-administration under FR1 and FR3 schedules of reinforcement. Finally, we examined the reinforcing efficacy of oxycodone in progressive-ratio (PR) probe tests. To determine whether any observed stress effects were specific to opioid reinforcement, we also evaluated the effects of adolescent stressor exposure on sucrose-maintained responding in separate animals.

## 2. Materials & Methods

### 2.1 Animals

Subjects were adolescent (PND 39, 125-150 g upon arrival) male and female Sprague Dawley rats acquired from Inotiv (Indianapolis, IN). Animals were initially pair-housed in same-sex pairs upon arrival under a 12-hour reverse light-dark cycle (lights off at 09:00am) and allowed to acclimate for a minimum of 5 days before beginning experimental manipulations.

Animals were provided enrichment and *ad libitum* access to food and water in the home cage throughout the study. All procedures were conducted in accordance with the NIH Guide for the Care and Use of Laboratory Animals and were approved by the Rowan University Institutional Animal Care and Use Committee. Behavioral studies took place between 10:00am and 4:00pm.

### 2.2 Acute adolescent stress & surgery

Within 5 to 9 days after arrival (PND 44-48), animals were exposed to a single 15 min session of concurrent physical restraint stress and predator odor exposure as described previously (Borodovitsyna et al., 2018). Briefly, animals were transported to a laboratory outside of the vivarium and placed in a rodent restrainer (Harvard Apparatus, Holliston, MA) within a chemical fume hood lined with disposable absorbent padding. A light-duty tissue wipe (11.4 x 21.0 cm) saturated with 25 µl of the fox-derived odor 2,4,5-trimethylthiazole (TMT; Sigma–Aldrich, St. Louis, MO) was placed directly in front of the restraint tube near the nose of the subject. 15 min later, animals were promptly removed from the restrainer and placed into a clean cage prepared with fresh bedding, food, and water. Animals were returned to the housing room 30 min later to mitigate the risk of TMT exposure to other animals in the vivarium. Control subjects were gently handled by an experimenter for 15 min within the housing room and then placed into a clean cage. Following stressor/control manipulations, all animals were singly housed for the remainder of the study.

The day immediately following stressor exposure or control handling (PND 45-49), animals destined for oxycodone self-administration were surgically prepared with chronic indwelling IV catheters under inhaled isoflurane anesthesia as previously described (Hinds et al., 2023; Manvich et al., 2016). Briefly, a custom-made silastic catheter (P1 Technologies, Roanoke, VA) was inserted into the right jugular vein and secured within the vessel via nonabsorbable sutures. The catheter was passed subcutaneously to an infusion port positioned between the scapulae from which an infusion cannula extended above the skin. Beginning on the day of surgery, catheters were flushed 5-6 days per week with 0.05 ml gentamicin (4 mg/ml) and locked with 0.1 ml of heparin (300 units/ml). When not in use, catheters were protected with a silastic obturator and a stainless-steel dust cap. For the first two days post-surgery, animals were administered the analgesic carprofen (2 mg, PO). Animals were allowed 6-7 days of recovery before the onset of oxycodone self-administration sessions.

### 2.3 IV Oxycodone Self-Administration

Oxycodone self-administration was conducted 5-6 days/week in operant chambers housed within ventilated, sound-attenuating cubicles (Med Associates, Fairfax, VT). Each session began with the extension of two levers into the operant chamber and illumination of a house light located on the opposite wall. An automated syringe pump (PHM100; Med Associates) located outside the cubicle held a 5cc syringe connected via Tygon tubing to a 22-g liquid swivel (Instech Laboratories, Plymouth Meeting, PA, USA) that was secured by a counterweighted arm above the operant chamber. A metal tether (Med Associates) protected an additional segment of Tygon tubing that extended from the output of the liquid swivel to the vascular access port of the subject. Med-PC V software (Med Associates) interfaced with each operant chamber to control outputs and record responses. Subject weights were entered into the self-administration program daily to ensure accuracy of oxycodone unit doses.

During self-administration sessions, successful completion of the ratio requirement on the active lever resulted in an infusion of oxycodone (∼ 2-4 s infusion duration) paired with illumination of the cue light above the active lever and termination of the house light for 20 s (timeout), during which active lever responses were recorded but were not reinforced. Inactive lever presses were also recorded throughout the session but had no consequences. Animals were provided a wooden chew block during self-administration sessions to mitigate damage from chewing of operant chamber components, a behavioral output which we and others have previously reported during oxycodone self-administration (Hinds et al., 2023) (Zanni et al., 2020). Sessions were terminated if animals earned 60 infusions or if 2 h elapsed, whichever occurred first. If animals earned 60 infusions, the levers were retracted and cue/house lights were turned off, but the animal remained in the operant chamber for the remainder of the 2 h session.

### 2.4 Acquisition and maintenance of oxycodone self-administration

A full overview of the oxycodone self-administration experimental timeline is depicted in **Fig. 1**. Beginning 6-7 days after stress exposure or control handling (PND 51-56), subjects (n=8 stress males, 9 control males, 10 stress females, 12 control females) were allowed to self-administer oxycodone (0.03 mg/kg/inf, IV) under a FR1 schedule of reinforcement for a maximum of 10 sessions. Animals were deemed to have acquired oxycodone self-administration if they earned ≥15 infusions in two consecutive sessions. If animals did not exhibit selectivity for the active lever at the time of acquisition (defined as emitting ≥75% total responses on the active lever), they continued on the FR1 schedule until attaining ≥75% active lever selectivity (5 females and 2 males required an additional 1-4 self-administration sessions following acquisition). Once self-administration was acquired and responding was selective for the active lever, each rat continued to self-administer oxycodone at FR1 for 3 sessions, followed by 7 sessions at FR3.

**Figure 1.**
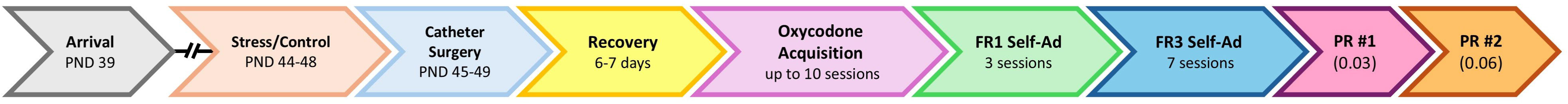
Oxycodone self-administration timeline. Following adolescent stress or control handling, rats were implanted with an intrajugular catheter and allowed to acquire self-administration of IV oxycodone (0.03 mg/kg/inf) for a maximum of 10 sessions. Once acquired and selective for the active lever, rats continued to a maintenance phase consisting of 3 sessions under FR1 and 7 sessions under FR3 schedules of reinforcement. Two PR test sessions were then performed in consecutive days with 0.03 mg/kg/inf (test 1) and 0.06 mg/kg/inf (test 2) IV oxycodone available as the reinforcer.

### 2.5 Progressive Ratio (PR)

On the two days immediately following the last FR3 session, animals underwent two consecutive test sessions using a PR schedule of reinforcement (one test per day; **Fig. 1**).

Responding was reinforced with 0.03 mg/kg/inf oxycodone during the first PR test and 0.06 mg/kg/inf oxycodone during the second PR test. During PR test sessions, the number of active lever responses required to receive a subsequent infusion was progressively increased as reported previously (1, 2, 4, 6, 9, 12, 15, 20, 25, 32, 40, 50, 62, 77, 95, 118, 145, 178, 219, 268, 328, 402, 492, 603… (Jordan et al., 2019; Richardson and Roberts, 1996). PR test sessions ended if 4 h elapsed or if 1 h passed without an infusion earned, whichever occurred first. The primary dependent measure was the “breakpoint”, defined as the last successfully completed ratio.

### 2.6 Sucrose Self-Administration

Separate groups of rats (n=8 stress males, 8 control males, 8 stress females, 8 control females) underwent sucrose pellet self-administration. Control/stressor exposure occurred on PND 45-46 as described above, with the exception that animals destined for sucrose pellet self-administration did not undergo catheter implantation. One week following stress/control handling, subjects were allowed to lever press for sucrose pellet reinforcement (45 mg, F0023 dustless precision pellet, Bio-Serv, Flemington, NJ) under a FR1 schedule of reinforcement.

Parameters for sucrose self-administration sessions were identical to those described for IV oxycodone self-administration sessions with the following exceptions: each session lasted 1 h, the maximum number of sucrose pellet reinforcers available was unrestricted, and the post-reinforcer timeout was 5 s. Following acquisition (defined as ≥ 15 reinforcers received in 2 consecutive sessions), animals continued to respond at FR1 for 3 sessions followed by 7 sessions at FR3. The day immediately following the last FR3 session, animals underwent one sucrose PR test under identical conditions described above for the oxycodone self-administering animals, with the exception that animals were reinforced with a single sucrose pellet.

### 2.7 Drugs

Oxycodone hydrochloride was provided by the National Institute on Drug Abuse Drug Supply Program (Research Triangle Park, NC) and dissolved at 0.18 mg/ml in sterile bacteriostatic saline. Oxycodone solutions were passed through a 0.22 μm filter and stored in sterile vials or syringes at 4°C between self-administration sessions. All drug doses are reported as the salt weight.

### 2.8 Statistical Analyses

#### Acquisition

Number of days to satisfy acquisition criteria for oxycodone or sucrose self-administration were analyzed using the Mantel-Cox (Log-Rank) test. Sex differences were assessed first, with stress and control animals collapsed within each sex. When no overall sex difference was detected, males and females were combined within each experimental group to compare acquisition in stress vs. control subjects. When sex differences were detected, Mantel-Cox analyses comparing stress vs. control subjects were performed separately in males and females.

#### FR1 and FR3 self-administration

Four female rats in the oxycodone self-administration study were unable to complete this phase of the experiment due to loss of catheter patency during these sessions (n=1 control) or unrelated health concerns (n=2 control, n=1 stress) and were therefore excluded from analysis of FR1-FR3 sessions and subsequent PR tests. Three-way mixed factors analysis of variance (ANOVA) with repeated measures on session and independent measures on experimental group (stress, control) and sex performed first for number of active and inactive lever presses, active and inactive lever response rates, and infusions earned. If the three-way ANOVA did not detect a main effect of sex nor any sex-dependent interactions, experimental groups were collapsed across sex and subsequently analyzed via two-way mixed factors ANOVA with repeated measures on session and independent measures on experimental group. Post-hoc pairwise comparisons were assessed using the Holm-Šídák test. Separate ANOVAs were applied to FR1 sessions and FR3 sessions because the expected reduction in operant responding during the transition to FR3 would have confounded singular analysis of the entire FR1-FR3 self-administration phase.

#### Progressive-ratio tests

Two female subjects in the oxycodone self-administration experiment were unable to complete both PR tests due to loss of catheter patency (n=1 stress) or unrelated health concerns (n=1 control) and were therefore excluded from the analysis. Separate unpaired t-tests were performed for each oxycodone test session (0.03 and 0.06 mg/kg/inf) and for the single sucrose PR test session to compare PR breakpoints. Sex differences were assessed first, and when no sex difference was detected, males and females were combined within each experimental group for subsequent analyses.

#### Statistical analysis software

Three-Way ANOVAs were performed using IBM SPSS Statistics software v29.0.2.0 (IBM, Armonk, NY). All other analyses were performed and figures were plotted using GraphPad Prism v10.4.1 (GraphPad Software, San Diego, CA). Significance was set at α = 0.05 for all statistical tests.

## 3. Results

### 3.1 Adolescent stress does not impact acquisition of oxycodone self-administration

All animals (39/39) acquired oxycodone self-administration within 10 sessions. There was no effect of sex on rate of acquisition (**Supplementary** Fig. 1) therefore male and female subjects in each experimental group were combined. No significant differences were observed in the rate of acquisition between stress and control groups (Mantel-Cox log-rank test; χ^2^_(1)_ = 1.23, p = 0.268; **Fig. 2**).

**Figure 2.**
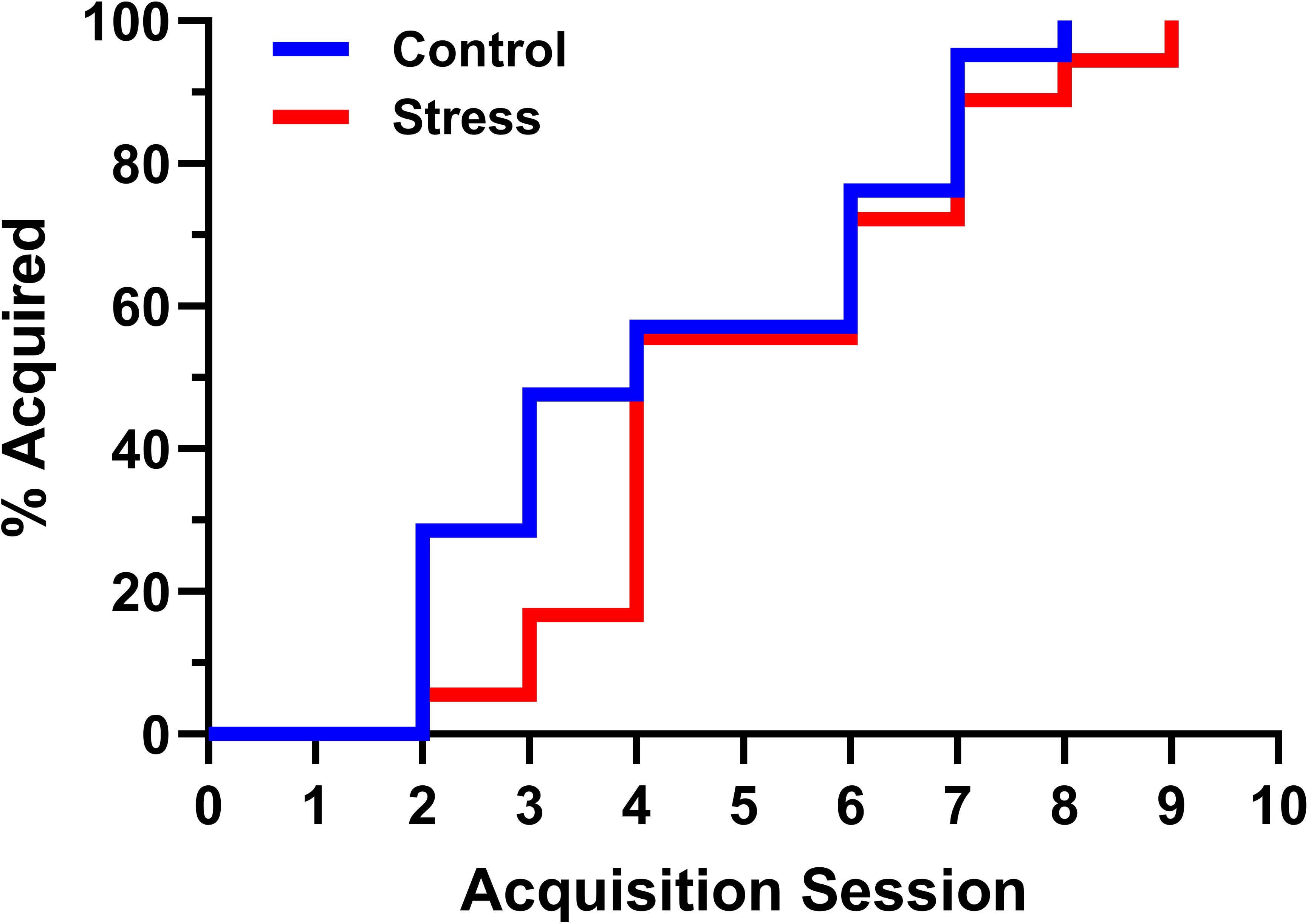
Rate of acquisition for IV oxycodone self-administration in stress and control rats. Kaplan-Meier reverse survival curves depicting the percentage of stress rats (n = 18, 8M 10F; red) and control rats (n = 21, 9M 12F; blue) that satisfied acquisition criteria for IV oxycodone self-administration over 10 sessions under a FR1 schedule of reinforcement.

### 3.2 No sex differences in oxycodone self-administration during FR1 or FR3 sessions

Three-way ANOVAs found no main effect of sex nor any sex-dependent interactions on active lever presses, active lever response rate, or infusions earned during FR1 sessions (**Supplementary Table 1**) or FR3 sessions (**Supplementary Table 2**). Therefore, males and females in each experimental group were combined for subsequent analyses.

### 3.3 FR1 sessions – oxycodone self-administration escalates in adolescent stressed rats

There was no effect of group on the number of active lever responses, however there was a significant main effect of session (F_(1.63, 53.62)_ = 4.38, p = 0.024) and a significant session × group interaction (F_(2, 66)_ = 3.19, p = 0.047), but post hoc comparisons did not detect differences in active lever responses across FR1 sessions in either group (**Fig. 3A**). When the number of active lever responses emitted was transformed to response rate by dividing the number of active lever presses by the total session time (which corrects for early session termination due to maximum reinforcer delivery), there was again a main effect of session (F_(1.49, 49.01)_ = 4.14, p = 0.032) but no main effect of group, and the session × group interaction narrowly missed significance (F_(2, 66)_ = 2.97, p = 0.058) (**Fig. 3B**). There was also no main effect of group on the number of infusions earned, however there was a main effect of session (F_(2, 66)_ = 12.23, p < 0.0001) and a significant session × group interaction (F_(2, 66)_ = 5.18, p = 0.008) (**Fig. 3C**). Post-hoc analyses revealed escalating intake in the stressed rats but not in control rats (**Fig. 3C**). There were no significant effects or interactions for inactive lever presses (**Fig. 3A**) and inactive lever response rate (**Fig. 3B**).

**Figure 3.**
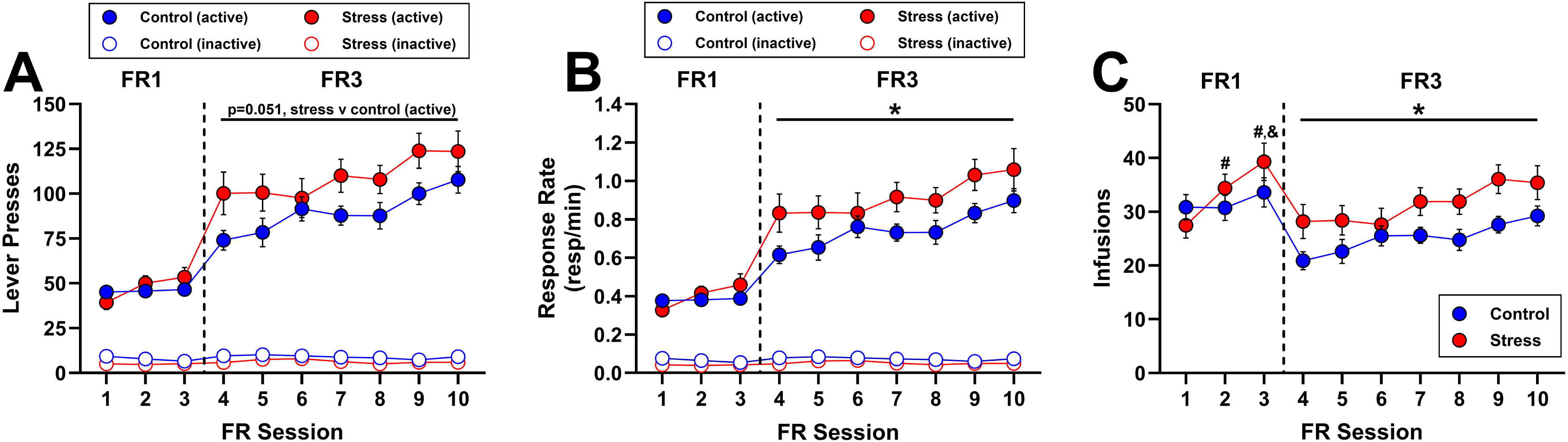
Adolescent stress-induced enhancement of IV oxycodone self-administration. Maintenance phase of IV oxycodone self-administration following acquisition in rats previously exposed to acute adolescent stress (n = 17, 8M 9F) or control handling (n = 18, 9M 9F). The maintenance phase consisted of 3 FR1 sessions (sessions 1-3) followed by 7 FR3 sessions (sessions 4-10). **(A)** Number of active and inactive lever responses. **(B)** Rates of responding on active and inactive levers. **(C)** Number of oxycodone infusions earned. Dashed lines indicate the transition between FR1 and FR3 schedules of reinforcement. All data are presented as mean ± SEM values. *p<0.05, main effect of group during FR3 sessions. ^#^p<0.05, compared to stress session 1. ^&^p<0.05, compared to stress session 2. Absence of error bars indicates that SEM values did not extend beyond the limits of the depicted symbol.

### 3.4 FR3 sessions – enhanced oxycodone self-administration in adolescent stressed rats

There was a trend towards a main effect of group on the number active lever responses during FR3 sessions (F_(1, 33)_ = 4.11, p = 0.051) and a significant main effect of session (F_(3.46, 114.0)_ = 7.48, p < 0.0001) but no significant session × group interaction (**Fig. 3A**). When active lever presses were transformed to response rate, there were significant main effects of group (F_(1, 33)_ = 4.21, p = 0.048) and session (F_(3.22, 106.2)_ = 7.09, p = 0.0002) with no significant session × group interaction, suggesting that oxycodone maintained higher rates of active lever responding in stressed rats stably throughout the FR3 phase (**Fig. 3B**). There was a similar main effect of group on the number of oxycodone infusions earned (F_(1, 33)_ = 5.29, p = 0.028) and a main effect of session (F_(3.35, 110.4)_ = 7.95, p < 0.0001), with no significant session × group interaction (**Fig. 3C**). There were no significant effects or interactions for inactive lever presses (**Fig. 3A**) and inactive lever response rate (**Fig. 3B**).

### 3.5 No effect of adolescent stress on PR responding for oxycodone

There were no sex differences in PR breakpoints for either dose of oxycodone (**Supplementary** Fig. 2), therefore males and females were combined within each experimental group. There was no significant effect of adolescent stress on PR breakpoint (**Fig. 4**) at either 0.03 mg/kg/inf (t_(31)_ = 0.48, p = 0.633) or 0.06 mg/kg/inf (t_(31)_ = 1.52, p = 0.139).

**Figure 4.**
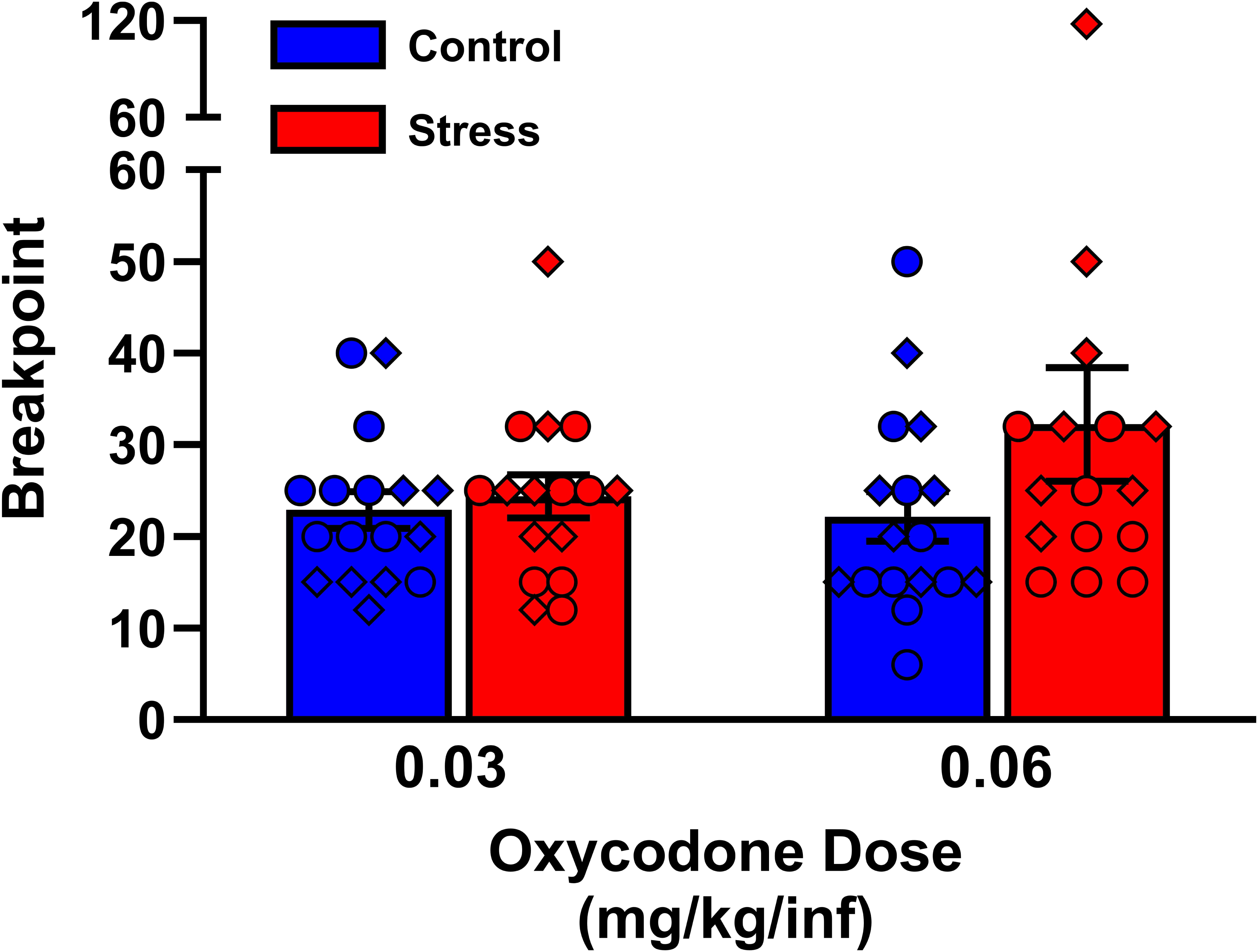
PR breakpoints for IV oxycodone in stress and control rats. IV oxycodone self-administration was assessed in two consecutive PR tests (0.03 and 0.06 mg/kg/inf, respectively) in rats previously exposed to acute adolescent stress (n = 16, 8M 8F; red) or control handling (n = 17, 9M 8F; blue) following acquisition and maintenance of oxycodone self-administration. Data are presented as individual data points (males, circles; females, diamonds) superimposed over mean ± SEM values.

### 3.6 Adolescent stress does not impact acquisition of sucrose pellet reinforcement

31/32 animals acquired sucrose pellet-maintained responding within 10 sessions. One rat (stress male) that failed to acquire was included in analysis of acquisition but excluded from the remainder of the study. Females acquired sucrose pellet self-administration faster than males (χ^2^_(1)_=4.51, p = 0.034; **Fig. 5A**), therefore the impact of adolescent stress on acquisition of sucrose reinforcement was examined separately in males and females. There was no effect of adolescent stress on acquisition in males (χ^2^_(1)_ = 0.33, p = 0.568; **Fig. 5B**) or females (χ^2^_(1)_ = 1.56, p = 0.212; **Fig. 5C**).

**Figure 5.**
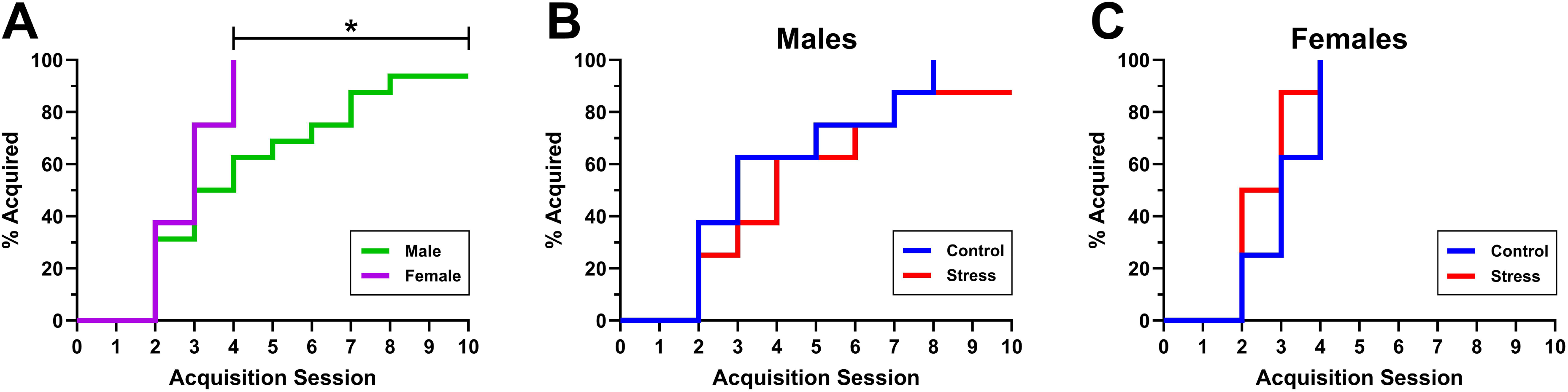
Rates of acquisition for sucrose-maintained responding. Kaplan-Meier reverse survival curves depicting the percentage of rats (n = 32, 16 M 16F) that satisfied acquisition criteria for sucrose pellet self-administration over 10 sessions under a FR1 schedule of reinforcement. Data are shown first separated by sex **(A)** and then subsequently subdivided by sex to compare stress vs. control groups in males **(B)** and females **(C)**. n = 8 per sex per group.

### 3.7 No sex differences in sucrose self-administration during FR1 or FR3 sessions

Three-way ANOVAs found no main effect of sex nor any sex-dependent interactions on active lever presses, active lever response rate, or number of pellets earned during FR1 sessions (**Supplementary Table 3**) or FR3 sessions (**Supplementary Table 4**). Males and females were therefore combined in each experimental group for subsequent analyses.

### 3.8 No effect of adolescent stress on sucrose self-administration during FR1 or FR3 sessions

For the FR1 phase, there were no main effects or interactions for active or inactive lever presses (**Fig. 6A**), active or inactive lever response rates (**Fig. 6B**), and number of sucrose pellets earned (**Fig. 6c**). For the FR3 phase, a main effect of session was detected for active lever responses emitted (F_(2.97, 86.05)_ = 5.59, p = 0.002), active lever response rate (F_(2.968, 86.08)_ = 5.60, p = 0.002), and number of sucrose pellets earned (F_(3.314, 96.12)_ = 4.34, p = 0.005). No other main effects or interactions were detected for any measures. These results suggest that sucrose-maintained responding and intake increased across the FR3 phase independent of adolescent stress history.

**Figure 6.**
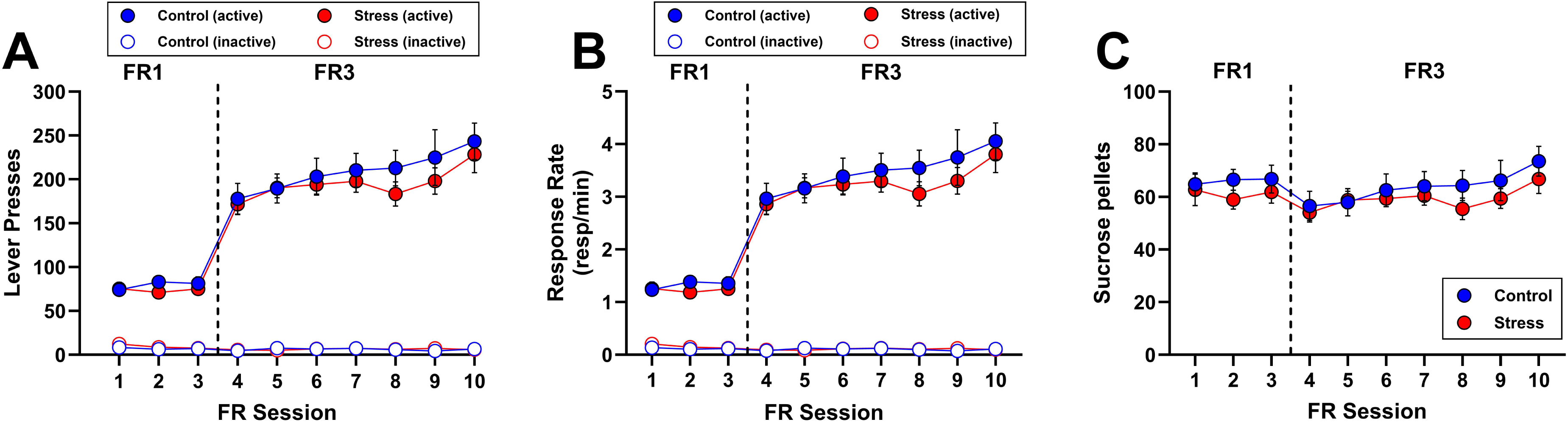
No effect of adolescent stress on sucrose pellet self-administration. Maintenance phase of sucrose pellet self-administration following acquisition in rats previously exposed to acute adolescent stress (n = 15, 7M 8F) or control handling (n = 16, 8M 8F). Other figure details are identical to those described in Fig. 3 with the exception that panel **C** here depicts numbers of sucrose pellets earned rather than IV oxycodone infusions.

### 3.9 No effect of adolescent stress on PR responding for sucrose

There was no sex difference in PR breakpoints for sucrose pellets (**Supplementary** Fig. 3), thus males and females were combined within each experimental group. There was no effect of adolescent stress on PR breakpoints for sucrose (t_(29)_ = 0.27, p = 0.791; **Fig. 7**).

**Figure 7.**
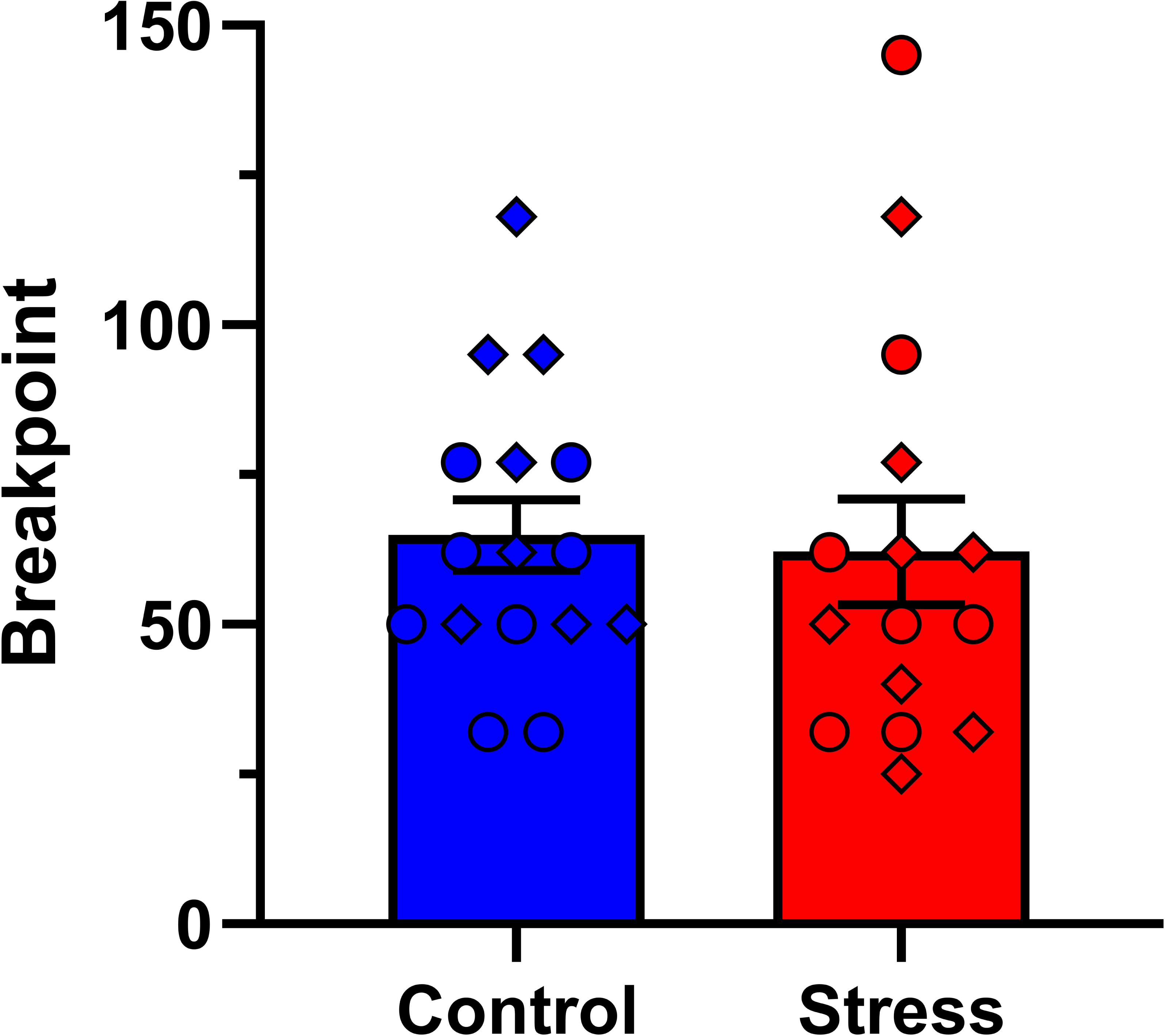
PR breakpoints for sucrose pellets in stress and control rats. Male (circles) and female (diamonds) rats previously exposed to adolescent stress (n = 15, 7M 8F; red) or control handling (n = 16, 8M 8F; blue) underwent a single sucrose PR test following acquisition and maintenance of sucrose self-administration. n = 16 control (8M, 8F), n=15 stress (7M, 8F). Data are presented as individual data points superimposed over mean ± SEM values.

## 4. Discussion

The present study investigated whether acute adolescent stress impacts IV oxycodone reinforcement in male and female rats during initial and later phases of drug intake. The major findings were that acute exposure to concurrent physical restraint and predator odor during adolescence did not affect the rate at which rats acquired IV oxycodone self-administration, but enhanced self-administration under FR schedules of reinforcement after acquisition. No sex differences were observed on any measure of oxycodone-reinforced responding, and we found no effect of acute adolescent stress on sucrose-reinforced responding, suggesting that the observed impacts of adolescent stress on oxycodone self-administration are not generalizable to nondrug rewards. These findings add to the preclinical literature describing stress-induced potentiation of the reinforcing effects of opioids and are in agreement with epidemiological data linking adolescent stress exposure to increased vulnerability for opioid misuse in humans (Afifi et al., 2012; Austin and Shanahan, 2018; Conroy et al., 2009; Dube et al., 2003; Swedo et al., 2020).

To our knowledge, the present study is the first to investigate the impact of adolescent stress on the initial reinforcing efficacy of oxycodone in a rodent self-administration model. Our finding that adolescent stress did not affect the rate at which rats acquired self-administration of 0.03 mg/kg/inf oxycodone is in agreement with two previous studies that reported no impact of adolescent stress on the acquisition of self-administration for remifentanil (Thorpe et al., 2020) or heroin (Singh et al., 2022). Collectively, these results suggest that adolescent stress does not appreciably modulate the initial reinforcing effects of opioids in rats, however given that all rats eventually acquired self-administration of 0.03 mg/kg/inf oxycodone in the present study, it is possible that use of a lower unit dose might have revealed subtler differences in initial sensitivity to oxycodone reinforcement between experimental groups (Carroll and Lac, 1997; Lynch et al., 2010).

Immediately following acquisition, we observed escalating oxycodone intake over the next three FR1 sessions selectively in the adolescent stressed animals. Because the self-administration behavior in the stressed group had not yet stabilized and may still have been increasing prior to the transition to the FR3 schedule, it is possible that extending the number of FR1 sessions may have allowed for an even further escalation and an unmasking of significant group differences during the FR1 phase. Indeed, when the schedule of reinforcement was subsequently increased to FR3, stressed rats exhibited greater overall levels of active lever responding and oxycodone intake as compared to their control counterparts, indicating that oxycodone functioned as a more effective reinforcer in the stressed animals during the FR3 phase. While we cannot rule out the possibility that greater oxycodone reinforcement in the stressed animals might also have been observed under the FR1 schedule had the animals been allowed to self-administer at FR1 for more sessions, it is also possible that the adolescent stress-induced enhancement of oxycodone self-administration is only evident under more demanding schedules of reinforcement. This would be consistent with a previous study reporting that footshock stress had no effect on initial acquisition of heroin self-administration in adult rats, but increased responding under a PR schedule (Shaham and Stewart, 1994), as well as several examples where experimental manipulations increased the reinforcing effects of psychostimulants specifically under demanding, but not continuous, schedules of reinforcement (Mendrek et al., 1998; Suto et al., 2002; Ward et al., 2003).

We tentatively interpret the observed increase in active lever responding and oxycodone infusions within the adolescent stress group as a stress-induced enhancement of oxycodone reinforcement, in line with interpretations by others where stress exposure similarly increased these measures in rodent models of drug intake (for review, see (Kamens et al., 2023)). However, we acknowledge that an increase in oxycodone-maintained responding might also be consistent with an attenuation of oxycodone effects and a rightward shift of the oxycodone dose-response function, especially since 0.03 mg/kg/inf oxycodone typically falls on the descending limb of the IV oxycodone dose-response curve under a FR schedule of reinforcement (Hinds et al., 2023; Mavrikaki et al., 2021, 2017). Dose-response studies or behavioral economics procedures applied to adolescent stress vs. control rats would be necessary to confirm whether the present shift in oxycodone reinforcement represents a potentiation (i.e., upward shift) or attenuation (i.e., rightward shift) of the reinforcing efficacy of 0.03 mg/kg/inf oxycodone. Notably however, a single study that examined the impact of early life/adolescent stress on self-administration of multiple doses of heroin reported an upward shift of the heroin dose-response curve (Singh et al., 2022), indicative of enhanced reinforcing efficacy which aligns with our interpretation of the present results.

Whereas animals exposed to acute adolescent stress exhibited greater oxycodone self-administration under the FR3 schedule of reinforcement, they did not emit greater levels of responding under a PR schedule. This result was unexpected as the PR test is generally considered to be a preferred method for quantitative assessment of changes in the reinforcing efficacy of drugs (Richardson and Roberts, 1996; Stafford et al., 1998). Moreover, stress-induced enhancement of opioid-maintained responding under PR schedules has previously been demonstrated with oral fentanyl (Shaham et al., 1993), oral oxycodone (Cornejo et al., 2025), and IV heroin (Shaham and Stewart, 1994), although it is noteworthy that these studies were all performed in adult animals and that stressor exposure occurred immediately prior to the PR test sessions. This methodology contrasts with our present study in which stressor exposure occurred on a single occasion during adolescence, several weeks prior to the eventual PR test sessions. It is therefore possible that the impacts of the single acute adolescent stressor in the present study were transient and might have been dissipating at the time of PR testing. It is also possible that other features of our study design affected the capacity to detect changes in PR breakpoints. For example, long term entrainment of responding under the PR schedule has been performed for self-administration of heroin (De Vries et al., 2003; Duvauchelle et al., 1998; Zhou et al., 2005) and morphine (De and Grasing, 2023), and it is possible that this approach might have revealed stress-dependent differences in breakpoints. Furthermore, we noted that doubling the unit dose of oxycodone during the second PR test from 0.03 to 0.06 mg/kg/inf yielded relatively similar breakpoint values, so it is possible that use of higher unit doses in the PR probe tests may be necessary to reveal differences between stress and control rats.

We found no effects of acute adolescent stress on sucrose-maintained responding during initial acquisition, maintenance under FR schedules, or a single PR probe test. This confirms that adolescent stress did not disrupt subjects’ ability to perform operant-conditioning tasks and suggests that the stress-induced enhancement of oxycodone self-administration is unlikely due to nonspecific alterations in reward seeking or motivated behavior. While there was no stress effect on sucrose-maintained responding, female rats displayed expedited acquisition as compared to males. Prior studies investigating acquisition of sucrose reinforcement in adolescent rats have produced conflicting results, with reports of either no sex difference in rate of acquisition (Lynch, 2008) or increased sucrose consumption in males during sucrose acquisition/training (rate of acquisition was not reported in this study) (Harmony et al., 2020). The reasons for these discrepant results are unclear, but there are notable procedural differences among the studies including testing during light vs. dark cycles, age of animals at onset of training, sucrose formulation (e.g., solution vs. pellet), and acquisition criteria employed.

While our study did not investigate the neurobiological mechanisms underlying stress-induced enhancement of oxycodone reward, it is possible that neuroadaptations within the locus coeruleus-norepinephrine (LC-NE) system may have at least partially contributed to the observed effects. As a central component to the brain’s stress response, the LC is inhibited by the release of endogenous opioids (Curtis et al., 2001; Valentino and Van Bockstaele, 2014, 2001).

Importantly, the acute adolescent stressor employed in this study induces a persistent anxiety-like phenotype and confers adaptations to LC physiology and opioid receptor expression in adolescent male rats that are associated with a reduced sensitivity of LC NE neurons to opioid-mediated inhibition (Borodovitsyna et al., 2022, 2018; Tkaczynski et al., 2022). It is intriguing to speculate that these LC-centric effects may in some part contribute to the adolescent stress-induced enhancement of oxycodone self-administration observed here, however future research using pharmacological, chemogenetic, optogenetic, and/or other manipulations targeting the LC-NE system will be necessary to characterize the potential contributions of LC-NE function to the stress-induced potentiation of oxycodone reinforcement observed here. It is also possible that neuroadaptations within mesocorticolimbic dopaminergic systems might contribute to the adolescent stress-induced potentiation of oxycodone self-administration, given that opioid-induced increases in dopaminergic neurotransmission are known to play a prominent role in their abuse-related properties (Di Chiara and North, 1992; Pierce and Kumaresan, 2006). For example, previous studies have demonstrated that adolescent social isolation/instability stress alters dopaminergic and/or glutamatergic signaling (Deutschmann et al., 2022; Karkhanis et al., 2019, 2016; Novoa et al., 2021), neuroactivational and neurochemical responses to drugs of abuse (Burke et al., 2013, 2010; Fosnocht et al., 2019; Howes et al., 2000; Karkhanis et al., 2015, 2014; Singh et al., 2022), and the expression of dopamine-and opioid-related genes including tyrosine hydroxylase, dopamine receptors, opioid peptides, and opioid receptors (Amancio-Belmont et al., 2020; Granholm et al., 2015; Karkhanis et al., 2019, 2016; Leonetti et al., 2025; Rodriguez-Arias et al., 2016) across various nodes of the mesocorticolimbic system. It is plausible that these alterations may influence vulnerability to the reinforcing properties of oxycodone, however the impacts of the restraint+TMT stressor utilized in the present study on mesocorticolimbic dopamine and opioid system dynamics have not yet been characterized and thus represent an important future direction for research.

## 5. Conclusions

The present study investigated the impact of acute adolescent stress on IV oxycodone self-administration in male and female rats. Although stressed rats acquired oxycodone self-administration at a similar rate to their control counterparts, they displayed escalating oxycodone intake during early FR1 sessions and enhanced oxycodone self-administration in subsequent FR3 sessions. These stress-induced impacts on oxycodone self-administration were not sex-dependent and did not extend to the nondrug reinforcer sucrose. To our knowledge, this is the first demonstration of adolescent stress-induced enhancement of IV oxycodone self-administration in rodents. While the neurobiological mechanisms underlying this phenomenon remain unknown, we have established a model which may be used to study the neural substrates and neuroadaptations contributing to this effect.

## CREDIT AUTHORSHIP CONTRIBUTION STATEMENT

**Corinne A. Gallagher:** Formal Analysis, Investigation, Data curation, Writing – Original Draft, Writing – Review & Editing, Visualization. **Daniel J. Chandler:** Conceptualization, Methodology, Resources, Funding Acquisition. **Daniel F. Manvich:** Conceptualization, Methodology, Software, Formal Analysis, Resources, Writing – Review & Editing, Visualization, Supervision, Project Administration, Funding Acquisition.

## FUNDING SOURCES

This work was supported by the National Institutes of Health/National Institute on Drug Abuse [R21DA062164 to DFM], [R21 DA052815 to DJC] and the Osteopathic Heritage Foundation [OHFE-F-2023-31 to DFM].

## DATA STATEMENT

The datasets generated and analyzed in the current study are available from the corresponding author upon reasonable request.

## DECLARATION OF INTEREST

Declarations of interest: none.

## Notes

### Competing Interest Statement

The authors have declared no competing interest.

