## Supplementary Material for "Effects of acute adolescent stress on the acquisition and maintenance of intravenous oxycodone self-administration in male and female rats"

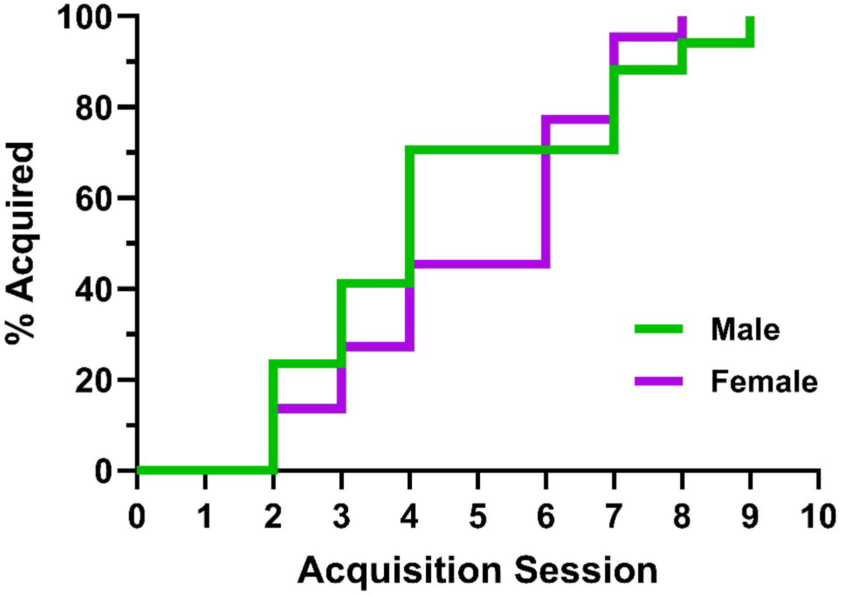

**Supplementary Figure 1. No sex difference in acquisition of IV oxycodone self-administration.** Shown are Kaplan-Meier reverse survival curves depicting the percentage of male (n = 17, green) and female (n = 22, purple) rats that satisfied acquisition criteria over 10 sessions under a FR1 schedule of reinforcement. Mantel-Cox log-rank analysis revealed no sex difference in the rate of acquisition ($\chi^2_{(1)} = 0.004$, $p = 0.948$).

**Supplementary Table 1**. Three-way ANOVA results for IV oxycodone self-administration during FR1 sessions (n = 8 stress males, 9 control males, 9 stress females, 9 control females). Significant *p* values are bolded.

| Factor | $F_{(df1, df2)}$ | *p* value |
|---|---|---|
| ***Active Lever Presses-FR1*** | | |
| **Group** | $F_{(1. 31)} = 0.14$ | 0.715 |
| **Sex** | $F_{(1, 31)} = 0.10$ | 0.749 |
| **Session** | $F_{(1.54, 47.72)} = 4.37$ | **0.026*** |
| **Session x sex** | $F_{(1.54, 47.72)} = 1.00$ | 0.358 |
| **Session x group** | $F_{(1.54, 47.72)} = 3.21$ | 0.061 |
| **Group x sex** | $F_{(1, 31)} = 0.06$ | 0.815 |
| **Session x group x sex** | $F_{(1.54, 47.72)} = 0.60$ | 0.509 |
| ***Inactive Lever Presses-FR1*** | | |
| **Group** | $F_{(1, 31)} = 2.34$ | 0.136 |
| **Sex** | $F_{(1, 31)} = 0.00$ | 0.987 |
| **Session** | $F_{(1.70, 52.63)} = 0.91$ | 0.396 |
| **Session x sex** | $F_{(1.70, 52.63)} = 1.22$ | 0.299 |
| **Session x group** | $F_{(1.70, 52.63)} = 1.18$ | 0.309 |
| **Group x sex** | $F_{(1, 31)} = 0.04$ | 0.847 |
| **Session x group x sex** | $F_{(1.70, 52.63)} = 0.55$ | 0.554 |
| ***Active Lever Response Rate-FR1*** | | |
| **Group** | $F_{(1, 31)} = 0.20$ | 0.656 |
| **Sex** | $F_{(1, 31)} = 0.04$ | 0.838 |
| **Session** | $F_{(1.40, 43.49)} = 4.09$ | **0.036*** |
| **Session x sex** | $F_{(1.40, 43.49)} = 1.15$ | 0.310 |
| **Session x group** | $F_{(1.40, 43.49)} = 2.95$ | 0.080 |
| **Group x sex** | $F_{(1, 31)} = 0.11$ | 0.741 |
| **Session x group x sex** | $F_{(1.40, 43.49)} = 0.61$ | 0.490 |
| ***Inactive Lever Response Rate-FR1*** | | |
| **Group** | $F_{(1, 31)} = 2.36$ | 0.135 |
| **Sex** | $F_{(1, 31)} = 0.00$ | 0.972 |
| **Session** | $F_{(1.73, 53.75)} = 0.91$ | 0.396 |
| **Session x sex** | $F_{(1.73, 53.75)} = 1.38$ | 0.259 |
| **Session x group** | $F_{(1.73, 53.75)} = 1.00$ | 0.366 |
| **Group x sex** | $F_{(1, 31)} = 0.05$ | 0.823 |
| **Session x group x sex** | $F_{(1,73, 53.75)} = 0.58$ | 0.542 |
| ***Infusions Earned-FR1*** | | |
| **Group** | $F_{(1, 31)} = 0.41$ | 0.529 |
| **Sex** | $F_{(1, 31)} = 0.85$ | 0.364 |
| **Session** | $F_{(1.45, 44.87)} = 12.11$ | **<0.001*** ** |
| **Session x sex** | $F_{(1.45, 44.87)} = 0.29$ | 0.681 |
| **Session x group** | $F_{(1.45, 44.87)} = 5.22$ | **0.016*** |
| **Group x sex** | $F_{(1, 31)} = 0.39$ | 0.536 |
| **Session x group x sex** | $F_{(1.45, 44.87)} = 1.19$ | 0.301 |

**Supplementary Table 2**. Three-way ANOVA results for IV oxycodone self-administration during FR3 sessions (n = 8 stress males, 9 control males, 9 stress females, 9 control females). Significant *p* values are bolded.

| Factor | $F_{(df1, df2)}$ | *p* value |
|---|---|---|
| ***Active Lever Presses-FR3*** | | |
| **Group** | $F_{(1, 31)} = 3.80$ | 0.060 |
| **Sex** | $F_{(1, 31)} = 0.13$ | 0.727 |
| **Session** | $F_{(3.34, 103.57)} = 7.16$ | **<0.001*** ** |
| **Session x sex** | $F_{(3.34, 103.57)} = 0.40$ | 0.777 |
| **Session x group** | $F_{(3.34, 103.57)} = 0.71$ | 0.563 |
| **Group x sex** | $F_{(1, 31)} = 0.40$ | 0.530 |
| **Session x group x sex** | $F_{(3.34, 103.57)} = 0.36$ | 0.800 |
| ***Inactive Lever Presses-FR3*** | | |
| **Group** | $F_{(1, 31)} = 1.85$ | 0.184 |
| **Sex** | $F_{(1, 31)} = 0.57$ | 0.458 |
| **Session** | $F_{(4.03, 125.05)} =1.13$ | 0.348 |
| **Session x sex** | $F_{(4.03, 125.05)} = 1.26$ | 0.290 |
| **Session x group** | $F_{(4.03, 125.05)} = 0.34$ | 0.851 |
| **Group x sex** | $F_{(1, 31)} = 1.54$ | 0.223 |
| **Session x group x sex** | $F_{(4.03, 125.05)} = 4.00$ | **0.004** ** |
| ***Active Lever Response Rate-FR3*** | | |
| **Group** | $F_{(1, 31)} = 3.90$ | 0.057 |
| **Sex** | $F_{(1, 31)} = 0.18$ | 0.678 |
| **Session** | $F_{(3.11, 96.43)} = 6.76$ | **<0.001*** ** |
| **Session x sex** | $F_{(3.11, 96.43)} = 0.43$ | 0.739 |
| **Session x group** | $F_{(3.11, 96.43)} = 0.45$ | 0.726 |
| **Group x sex** | $F_{(1, 31)} = 0.49$ | 0.489 |
| **Session x group x sex** | $F_{(3.11, 96.43)} = 0.36$ | 0.788 |
| ***Inactive Lever Response Rate-FR3*** | | |
| **Group** | $F_{(1, 31)} = 1.79$ | 0.190 |
| **Sex** | $F_{(1, 31)} = 0.53$ | 0.471 |
| **Session** | $F_{(3.99, 123.59)} = 1.13$ | 0.347 |
| **Session x sex** | $F_{(3.99, 123.59)} = 1.23$ | 0.301 |
| **Session x group** | $F_{(3.99, 123.59)} = 0.33$ | 0.859 |
| **Group x sex** | $F_{(1, 31)} = 1.53$ | 0.226 |
| **Session x group x sex** | $F_{(3.99, 123.59)} = 3.83$ | **0.006** ** |
| ***Infusions Earned-FR3*** | | |
| **Group** | $F_{(1, 31)} = 4.92$ | **0.034*** |
| **Sex** | $F_{(1, 31)} = 0.17$ | 0.681 |
| **Session** | $F_{(3.23, 100.0)} = 7.63$ | **<0.001*** ** |
| **Session x sex** | $F_{(3.23, 100.0)} = 0.38$ | 0.780 |
| **Session x group** | $F_{(3.23, 100.0)} = 0.86$ | 0.469 |
| **Group x sex** | $F_{(1, 31)} = 0.02$ | 0.881 |
| **Session x group x sex** | $F_{(3.23, 100.0)} = 0.55$ | 0.662 |

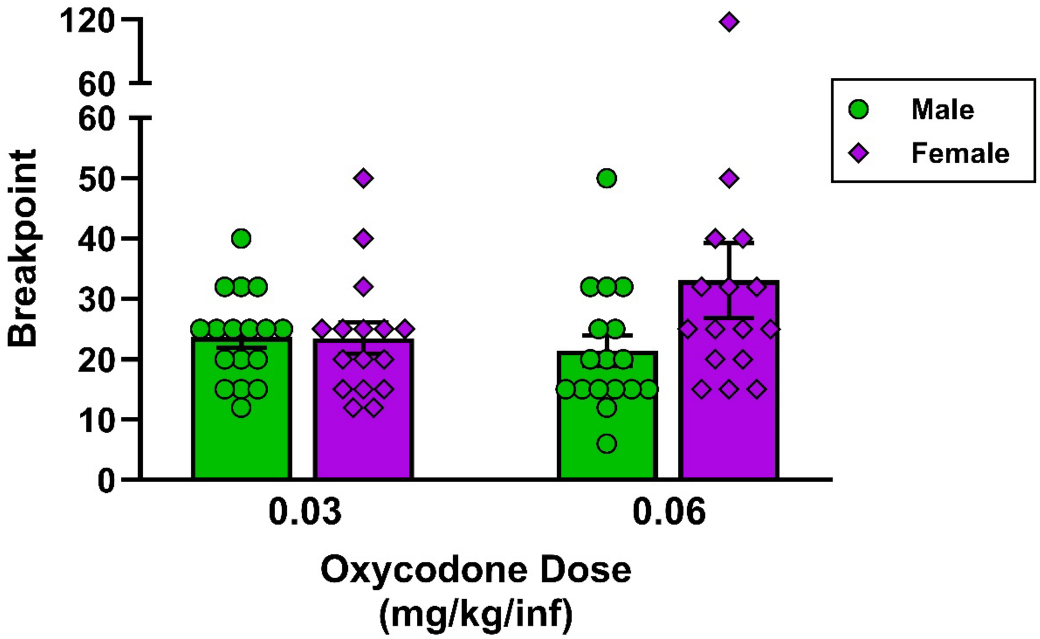

**Supplementary Figure 2. No sex difference in PR breakpoints for IV oxycodone at 0.03 or 0.06 mg/kg/inf.** n = 17 males, 16 females. Data are presented as individual data points superimposed over bars depicting mean ± SEM values. 0.03 mg/kg/inf, $t_{(31)} = 0.07$, $p = 0.948$; 0.06 mg/kg/inf, $t_{(31)} = 1.78$, $p = 0.085$.

**Supplementary Table 3.** Three-way ANOVA results for sucrose pellet self-administration during FR1 sessions (n = 7 stress males, 8 control males, 8 stress females, 8 control females). Significant *p* values are bolded.

| Factor | $F_{(df1, df2)}$ | *p* value |
|---|---|---|
| ***Active Lever Press-FR1*** | | |
| **Group** | $F_{(1, 27)} = 0.47$ | 0.499 |
| **Sex** | $F_{(1, 27)} = 0.39$ | 0.536 |
| **Session** | $F_{(1.64, 44.26)} = 0.52$ | 0.562 |
| **Session x sex** | $F_{(1.64, 44.26)} = 1.27$ | 0.287 |
| **Session x group** | $F_{(1.64, 44.26)} = 1.45$ | 0.245 |
| **Group x sex** | $F_{(1, 27)} = 0.80$ | 0.380 |
| **Session x group x sex** | $F_{(1.64, 44.26)} = 0.51$ | 0.569 |
| ***Inactive Lever Press-FR1*** | | |
| **Group** | $F_{(1, 27)} = 1.09$ | 0.305 |
| **Sex** | $F_{(1, 27)} = 7.82$ | **0.009**** |
| **Session** | $F_{(1.91, 51.64)} = 2.76$ | 0.075 |
| **Session x sex** | $F_{(1.91, 51.64)} = 1.91$ | 0.161 |
| **Session x group** | $F_{(1.91, 51.64)} = 0.80$ | 0.452 |
| **Group x sex** | $F_{(1, 27)} = 0.97$ | 0.334 |
| **Session x group x sex** | $F_{(1.91, 51.64)} = 2.21$ | 0.122 |
| ***Active Response Rate-FR1*** | | |
| **Group** | $F_{(1, 27)} = 0.48$ | 0.496 |
| **Sex** | $F_{(1, 27)} = 0.39$ | 0.536 |
| **Session** | $F_{(1.64, 44.29)} = 0.51$ | 0.568 |
| **Session x sex** | $F_{(1.64, 44.29)} = 1.27$ | 0.286 |
| **Session x group** | $F_{(1.64, 44.29)} = 1.44$ | 0.247 |
| **Group x sex** | $F_{(1, 27)} = 0.79$ | 0.383 |
| **Session x group x sex** | $F_{(1.64, 44.29)} = 0.50$ | 0.575 |
| ***Inactive Response Rate-FR1*** | | |
| **Group** | $F_{(1, 27)} = 1.12$ | 0.299 |
| **Sex** | $F_{(1, 27)} = 7.81$ | **0.009**** |
| **Session** | $F_{(1.91, 51.64)} = 2.76$ | 0.075 |
| **Session x sex** | $F_{(1.91, 51.64)} = 1.80$ | 0.176 |
| **Session x group** | $F_{(1.91, 51.64)} = 0.84$ | 0.435 |
| **Group x sex** | $F_{(1, 27)} = 0.93$ | 0.343 |
| **Session x group x sex** | $F_{(1.91, 51.64)} = 2.24$ | 0.119 |
| ***Reinforcers Earned-FR1*** | | |
| **Group** | $F_{(1, 27)} = 0.67$ | 0.421 |
| **Sex** | $F_{(1, 27)} = 0.40$ | 0.535 |
| **Session** | $F_{(1.69, 45.74)} = 0.15$ | 0.829 |
| **Session x sex** | $F_{(1.69, 45.74)} = 0.85$ | 0.419 |
| **Session x group** | $F_{(1.69, 45.74)} = 0.36$ | 0.662 |
| **Group x sex** | $F_{(1, 27)} = 1.27$ | 0.270 |
| **Session x group x sex** | $F_{(1.69, 45.74)} = 0.37$ | 0.659 |

**Supplemental Table 4.** Three-way ANOVA results for IV oxycodone self-administration during FR3 sessions (n = 7 stress males, 8 control males, 8 stress females, 8 control females). Significant *p* values are bolded.

| Factor | $F_{(df1, df2)}$ | *p* value |
|---|---|---|
| ***Active Lever Press-FR3*** | | |
| **Group** | $F_{(1, 27)} = 0.36$ | 0.554 |
| **Sex** | $F_{(1, 27)} = 0.06$ | 0.814 |
| **Session** | $F_{(3.14, 84.90)} = 5.88$ | **<0.001*** ** |
| **Session x sex** | $F_{(3.14, 84.90)} = 1.77$ | 0.156 |
| **Session x group** | $F_{(3.14, 84.90)} = 0.44$ | 0.735 |
| **Group x sex** | $F_{(1, 27)} = 0.83$ | 0.370 |
| **Session x group x sex** | $F_{(3.14, 84.90)} = 0.96$ | 0.421 |
| ***Inactive Lever Press-FR3*** | | |
| **Group** | $F_{(1, 27)} = 0.001$ | 0.973 |
| **Sex** | $F_{(1, 27)} = 3.63$ | 0.067 |
| **Session** | $F_{(4.32, 116.68)} = 0.75$ | 0.569 |
| **Session x sex** | $F_{(4.32, 116.68)} = 0.34$ | 0.867 |
| **Session x group** | $F_{(4.32, 116.68)} = 0.97$ | 0.431 |
| **Group x sex** | $F_{(1, 27)} = 0.64$ | 0.430 |
| **Session x group x sex** | $F_{(4.32, 116.68)} = 0.94$ | 0.449 |
| ***Active Response Rate-FR3*** | | |
| **Group** | $F_{(1, 27)} = 0.36$ | 0.554 |
| **Sex** | $F_{(1, 27)} = 0.06$ | 0.814 |
| **Session** | $F_{(3.15, 84.93)} = 5.90$ | **<0.001*** ** |
| **Session x sex** | $F_{(3.15, 84.93)} = 1.78$ | 0.155 |
| **Session x group** | $F_{(3.15, 84.93)} = 0.44$ | 0.736 |
| **Group x sex** | $F_{(1, 27)} = 0.83$ | 0.370 |
| **Session x group x sex** | $F_{(3.15, 84.93)} = 0.96$ | 0.419 |
| ***Inactive Response Rate-FR3*** | | |
| **Group** | $F_{(1, 27)} = 0.002$ | 0.961 |
| **Sex** | $F_{(1, 27)} = 3.72$ | 0.065 |
| **Session** | $F_{(4.30, 116.05)} = 0.74$ | 0.573 |
| **Session x sex** | $F_{(4.30, 116.05)} = 0.32$ | 0.877 |
| **Session x group** | $F_{(4.30, 116.05)} = 1.04$ | 0.392 |
| **Group x sex** | $F_{(1, 27)} = 0.63$ | 0.433 |
| **Session x group x sex** | $F_{(4.30, 116.05)} = 0.94$ | 0.451 |
| ***Reinforcers Earned-FR3*** | | |
| **Group** | $F_{(1, 27)} = 0.47$ | 0.498 |
| **Sex** | $F_{(1, 27)} = 0.001$ | 0.970 |
| **Session** | $F_{(3.52, 94.94)} = 4.52$ | **0.003** ** |
| **Session x sex** | $F_{(3.52, 94.94)} = 1.66$ | 0.174 |
| **Session x group** | $F_{(3.52, 94.94)} = 0.52$ | 0.701 |
| **Group x sex** | $F_{(1, 27)} = 1.40$ | 0.246 |
| **Session x group x sex** | $F_{(3.52, 94.94)} = 0.70$ | 0.574 |

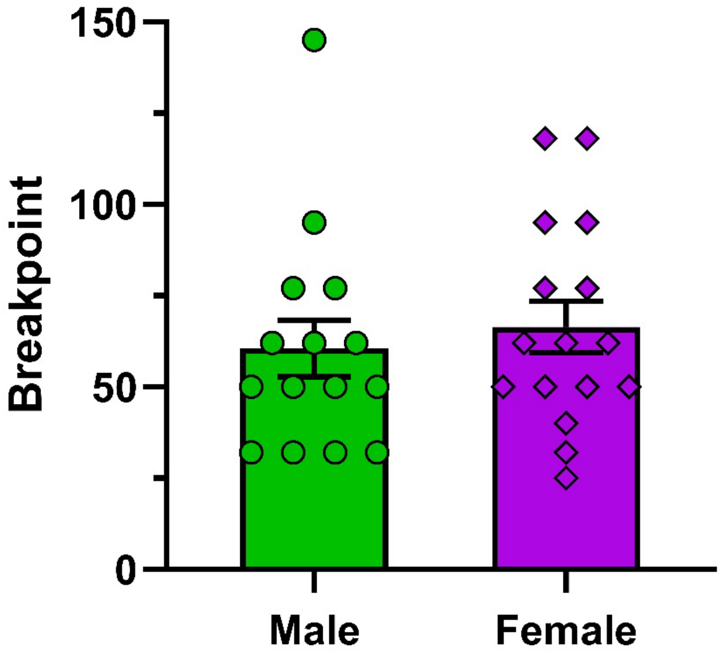

**Supplementary Figure 3. No sex difference in PR breakpoints for sucrose pellets.** n = 15 males, 16 females. Data are presented as individual data points superimposed over bars depicting mean ± SEM values. $t_{(29)} = 0.57$, $p = 0.576$.
